# Female ticks (*Ixodes scapularis*) infected with *Borrelia burgdorferi* have increased overwintering survival, with implications for tick population growth

**DOI:** 10.1101/2022.12.07.519462

**Authors:** Amal El Nabbout, Laura V. Ferguson, Atsushi Miyashita, Shelley A. Adamo

## Abstract

The tick, *Ixodes scapularis*, vectors pathogens such as *Borrelia burgdorferi*, the bacterium that causes Lyme Disease. Over the last few decades *I. scapularis* has expanded its range, introducing a novel health threat into these areas. Warming temperatures appear to be one cause of its range expansion to the north. However, other factors are also involved. We show that unfed adult female ticks infected with *B. burgdorferi* have greater overwintering survival than uninfected female ticks. Locally collected adult female ticks were placed in individual microcosms and allowed to overwinter in both forest and dune grass environments. In the spring we collected the ticks and tested both dead and living ticks for *B. burgdorferi* DNA. Infected ticks had greater overwintering survival compared with uninfected ticks every winter for three consecutive winters in both forest and dune grass environments. We discuss the most plausible explanations for this result. The increased winter survival of adult female ticks could enhance tick population growth. Our results suggest that, in addition to climate change, *B. burgdorferi* infection itself may be promoting the northern range expansion of *I. scapularis*. Our study highlights how pathogens could work synergistically with climate change to promote host range expansion.

## Introduction

Winter is a stressful time for arthropods. Low temperatures can cause tissue injury and death, limiting the northern ranges of many insects and their relatives, such as ticks (Williams et al. 2015). The medically-important tick, *Ixodes scapularis*, dies at temperatures below −10° to - 12°C (Burks et al. 1996; Vandyk et al. 1996), and even relatively mild winter temperatures (e.g.−3°C) may be lethal when humidity is high (i.e. due to inoculative freezing (Burks et al. 1996)). This cold intolerance may have historically excluded *I. scapularis* from northern regions of North America (Estrada-Peña 2002). Nevertheless, factors other than temperature play a role in determining overwintering survival (Lindsay et al. 1995; Lindsay et al. 1998; Brunner et al. 2014; Burtis et al. 2016; Burtis & Pflueger 2017; Clow et al. 2017; Kilpatrick et al. 2017; Hammond-Collins et al. 2022; Volk et al. 2022). For example, *I. scapularis* can successfully overwinter in regions that are below their thermal limits based on winter air temperatures (e.g. Northwestern Ontario (Lindsay et al. 1995), Upstate New York (Burtis et al. 2019), Maine (Volk et al. 2022)). Ticks can survive in these areas because they have adaptations, such as burrowing into leaf litter, that protect them from lethal temperatures (Brunner et al. 2012; Linske et al. 2019). However, overwintering survival is highly variable across years (Lindsay et al. 1998) (Volk et al. 2022), and our understanding of the various factors that determine overwintering success remains incomplete. We examine the possibility that infection with the bacterium *Borrelia burgdorferi* (the causative agent of Lyme Disease (Burgdorfer et al. 1982)) is also a factor in determining overwintering survival. We focus on the impact of infection on the overwintering survival of adult female ticks, because the overwintering survival of adult females has an impact on the size of the upcoming year’s tick population. Despite their importance, few studies (Lindsay et al. 1995; Lindsay et al. 1998) examine overwintering in adult females. We test whether infected adult female ticks have greater overwintering survival than non-infected females, and what this effect may mean for its northward expansion.

After hatching from eggs, *I. scapularis* larvae climb on vegetation and position themselves to find a host (i.e. quest) (for life cycle review see Eisen et al. 2012; Kilpatrick et al. 2017; Gray & Kahl 2022). Larvae attach themselves to a passing small mammal (e.g. mice). Mice such as the white footed mouse (*Peromyscus leucopus*) can tolerate high numbers of *B. burgdorferi* spirochetes in their blood when infected, allowing them to transmit *B. burgdorferi* to *I. scapularis* ticks that feed on them (Eisen et al. 2012). After a single feeding, the larvae molt into nymphs. *B. burgdorferi* persists in *I. scapularis* even after a molt (Helble et al. 2021; Pal et al. 2021). Questing nymphs feed on a range of mammals. Nymphs that were infected as larvae are able to transmit *B. burgdorferi* to its host (Gray & Kahl 2022). Uninfected nymphs can acquire *B. burgdorferi* if they feed on an infected mammal that is capable of transmitting it. After the second feeding, nymphs molt into adults. Adult females typically feed on larger mammals, such as a deer, on which they typically find a mate (Gray & Kahl 2022). After feeding, the female tick produces her eggs and then dies. Infected adult females are unlikely to find a mammalian host capable of transmitting *B. burgdorferi* to another tick (Halsey et al. 2018), preventing the bacteria from continuing their life cycle. Large mammals such as deer do not transmit *B. burgdorferi* to other ticks (Gray & Kahl 2022). However, although the *B. burgdorferi* bacteria found in adult females will have little or no reproductive success, the bacteria still interact physiologically with their host (Kurokawa et al. 2022).

Adults, nymphs and larvae can overwinter (Brunner et al. 2012). However, they have limited capacity for movement below 4°C (Duffy & Campbell 1994), making them unable to find a host or escape predators. Although the lack of questing (i.e. searching for a host) in winter means that they are without food for the season, it does allow them to remain within their insulated spaces and avoid the relatively lower ambient air temperatures (Burtis et al. 2019). Leaf litter provides insulation, resulting in warmer temperatures and higher relative humidity relative to the ambient air (Linske et al. 2019). However, overwintering survival can be low, even when ticks are provided with leaf litter (Lindsay et al. 1998). Overwintering survival of unfed adult females can vary from 27% to 78% depending on the year (Lindsay et al. 1998). If infection enhances overwintering survival, some of this variability may be related to the prevalence of infection in a population, that also varies from year to year (e.g. Scott et al. 2016).

Typically pathogens produce damage, reducing a host’s condition (Boucias & Penland, 1994). Such pathology would be expected to reduce overwintering survival. However, *B. burgdorferi* appears to have little or no negative impact on its host, although it does activate tick immune responses (Kurokawa et al. 2020). When microbes have little negative impact, they can enhance cold tolerance in arthropods (Ferguson et al. 2018). In the related tick *Ixodes ricinus*,nymphs infected with *B. burgdorferi* were more likely to survive stressful environmental conditions (e.g. low relative humidity (Herrmann & Gern 2010) and cold temperatures (Herrmann & Gern 2013) than uninfected ticks. How pathogens alter host cold tolerance in arthropods remains poorly understood (Overgaard & MacMillan 2017), but pathogens could improve the winter survival of their hosts in a few ways. Pathogens could actively manipulate hosts (e.g. ticks) to produce molecules that enhance host overwintering survival. This mechanism has been suggested as an explanation for the effect of *Borrelia* on *I. ricinis* (Herrmann & Gern, 2015). In *I. scapularis*, infection with the pathogen *Anaplasma phagocytophilum*, leads to the production of the protein IAFGP (*Ixodes* anti-freeze glycoprotein) (Neelakanta et al. 2010) (Heisig et al. 2014). This protein has both anti-microbial and cryoprotectant properties (Heisig et al. 2014). It enhances cold tolerance in the short-term under laboratory conditions (Neelakanta et al. 2010). Similarly, other molecules, such as heat shock proteins, are produced by organisms in response to stressors such as pathogens (Adamo 2022), and these molecules can protect organisms from a range of stressors (King & MacRae 2015), including low temperatures (Overgaard & MacMillan 2017), a phenomenon known as cross-tolerance (Sinclair et al. 2013). In this case, the pathogen is not manipulating the host, the host is activating an integrated defense response that includes broad protection from a number of stressors (Adamo 2022). *I. scapularis* infected with *B. burgdorferi* show increased gene expression for heat shock proteins (Hart et al. 2022), and other molecules (Narashimhan et al. 2017) known to increase cold tolerance in arthropods (Overgaard & MacMillan 2017). However, despite the existence of plausible mechanisms by which *B. burgdorferi* might enhance *I. scapularis* overwintering survival, it has not been demonstrated that infection with *B. burgdorferi* (or any other pathogen) increases overwintering success in *I. scapularis* under natural winter conditions. We examine whether there is an interaction between infection and winter survival in adult females.

If infected adult female ticks are more cold-tolerant, then this could increase their ability to survive the winter. More adult females in the population could translate into a rise in the number of larval ticks in the next generation. Enhanced growth of tick populations could also contribute to the northward expansion of *I. scapularis*.

## Materials and Methods

### Field Study

Field work was conducted at the Harrison Lewis Coastal Discovery Centre (HLC), Port L’Hebert, NS (HLC43°49’04.2” N 64°53’40.4” W), on the south shore of Nova Scotia. This area is experiencing increasing tick populations (Hatchette et al. 2015). The work was conducted over three winters (2018-2019, 2019-2020, and 2020-2021). Studies were approved by the University Committee on Laboratory Animals at Dalhousie University (protocol number: I18-06).

*Ixodes scapularis* were collected by dragging (Clow et al. 2017) from Thomas Raddall Provincial Park, located adjacent to the HLC. Following collection, ticks were placed in individual microcosms using a design modified from Burtis (Burtis 2017) as described in Adamo et al. (2022) and repeated here. Adult female ticks were placed in individual microcosms topped with maple/oak leaf litter (approximately 2 g). The microcosms were placed in the ground to the depth of the leaf litter (about 5 cm). Microcosms were placed in either: a) a coastal forest containing a mix of oak (*Quercus rubra*), maple (*Acer rubrum*) and spruce (*Picea rubens* and *Picea glauca*) or, b) within the dune grass next to a sand beach. Both areas harbour *I. scapularis* ticks in the spring, summer and fall. Microcosms were collected in the spring, and living and dead ticks were tested for the presence of *B. burgdorferi* DNA using qPCR.

The microcosm was an open plastic tube (15.24 cm in length x 1.27 cm in width), with mesh glued to each end, secured by a plastic cap. The plastic caps had a central hole that allowed rain and snow to enter in the top and connected the soil with the open bottom of the tube. The tubes contained sand and vermiculite (75% w/w sand and 25% w/w vermiculite) to a depth of 5 cm in the tube. The tubes were placed 5 cm into the ground. On top of the sand/vermiculite was 1 cm (approx. 2 g) of cut oak (*Quercus rubra* L.) and maple (*Acer rubrum* L.) leaves. A single tick was placed into each microcosm and then the microcosm was placed under a coarse netting (NuVue^®^ (78.7cm L x 78.7 cm W x 83.8 cm H). A data logger (HOBO MX2301A) was placed in each netting enclosure with a probe 5 cm underground between the microcosms; the logger was covered with a solar radiation shield (Onset RS3-B) and the logger automatically recorded temperature and relative humidity every hour. In 2018 (N=36), tick vials and data loggers were placed in in either dune grass or in a mixed maple/oak/spruce stand of trees (Fig. 1). In years 2 and 3 (Fig. 1), we placed six microcosms under a single netting rather than placing all microcosms for one study site under the same netting enclosure. Additionally, an extra study location was created for each site, both of which were also located at the HLC; as a result, there were two forest sites (forest 1 and forest 2) and two dune grass sites (dune grass 1 and dune grass 2) in years 2 and 3. In year 2, there were 145 ticks placed into individual microcosms. In year 3, there were 120 ticks placed into individual microcosms with 6 microcosms per netting enclosure.

**Figure 1.**
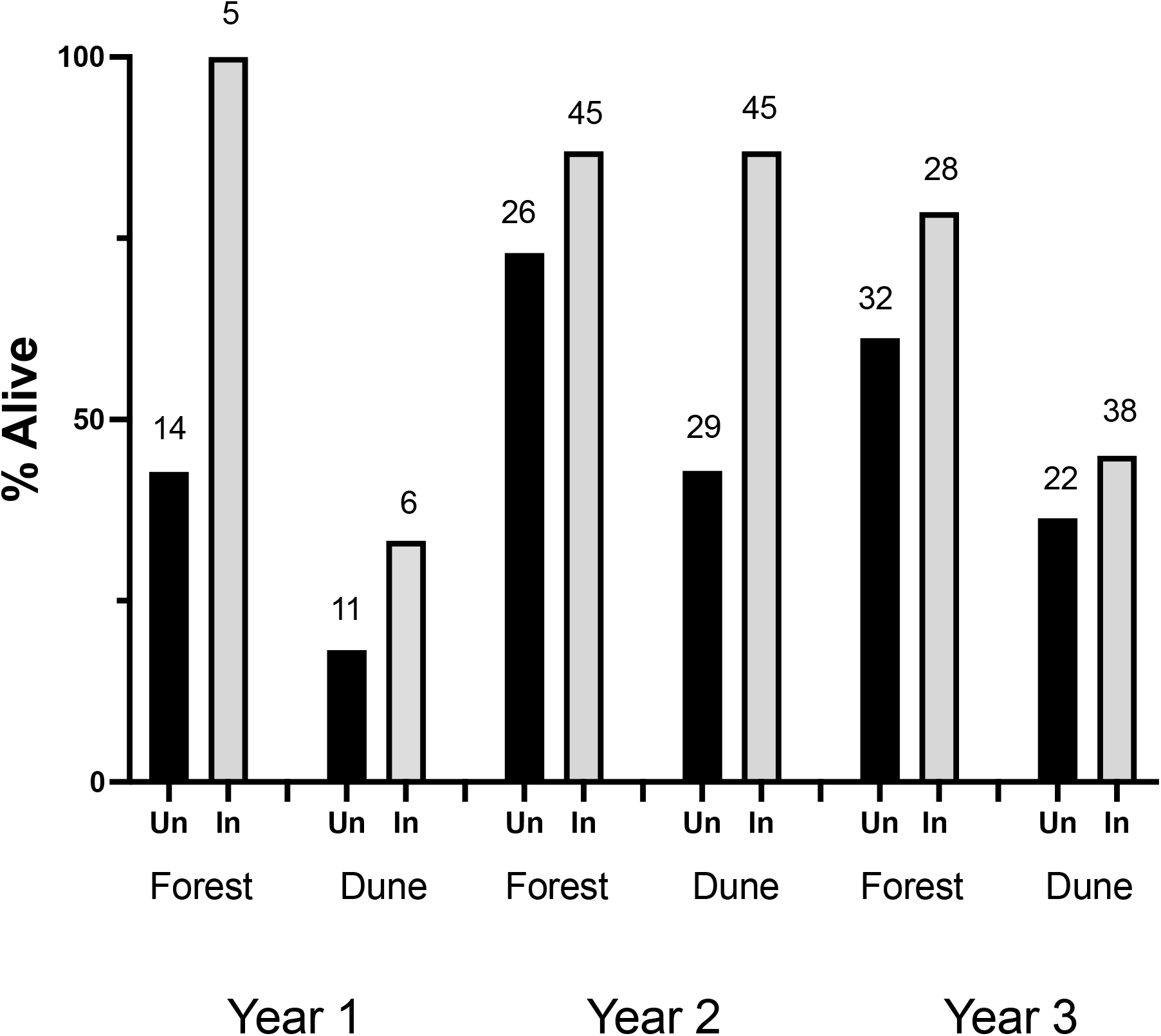
Percentage of *I. scapularis* infected with *B. burgdorferi* that survive the winter compared with uninfected ticks at each study site over the three winters. Numbers above each bar indicate the sample size. Dark bars denote uninfected ticks, while light bars denote infected ticks. Un=uninfected, In=Infected.

In Year 1 only, ticks were added to the microcosms as they were collected (from October 25, 2018 to Nov. 15, 2018). In years 2 and 3, ticks were held at 4° C and then all ticks were placed in microcosms in the field on the same day (Year 2, Nov 7, 2019; Year 3 Nov. 28, 2020). Ticks were retrieved on February 8, 2019 in Year 1. In year 2, ticks were retrieved on March 21, 2020, while in year 3 they were retrieved on March 23, 2021.

At the end of the field season, ticks were recovered by carefully sieving the tubes. Ticks that were immobile after collection were exposed to both a gentle exhalation by the experimenter (AEN) and probing with a paint brush. Ticks that showed no movement were tested again after 24 h at room temperature (approximately 22°C). If they still showed no movement they were deemed to be dead. Both dead and living ticks were tested for the presence of *B. burgdorferi* DNA using qPCR, following the procedure of Courtney et al. (2004), as described in Adamo et al. (2022) (see Supplementary Information). Each DNA sample was tested to ensure that the DNA was intact and not degraded. DNA integrity was assessed by gel electrophoresis (Aranda et al. 2012) using a Gel Doc™ XR+ Digital Gel Imaging System^®^ from Bio-Rad^®^. All of the DNA we extracted from dead ticks was intact showing distinct bands on the gel.

Because of the importance of demonstrating that we could accurately assess *Borrelia* DNA collected from dead ticks, we performed a lab study in which we simulated extreme winter conditions (i.e. daily freeze/thaw). We assessed the presence of *Borrelia* in one half of a sectioned adult female tick (n=30) immediately after cutting the tick in two, and the other half was assessed 10 weeks later. The second half of the tick was incubated in a fluctuating thermal regime incubator (−5°C to 10°C) in 12-hour increments for 10 weeks. We determined that we could accurately measure infection in ticks that had been dead for 10 weeks under these conditions (Supplementary Information, Table S2). There was 100% concordance between results at time 0 and 10 weeks later (Table S2).

### Statistical Analysis

Because of methodological differences across years, we analyzed the data using three 3-way Log-Linear analyses. Data from all 3 years fit the assumptions for a log-linear model. Post hoc tests were controlled for a false discovery rate using the Benjamini-Hochberg procedure (Benjamini & Hochberg 1995); www.biostathandbook.com/benjaminihochberg.xls). Tests were performed using SPSS (ver. 27) and Prism (GraphPad, ver. 9.4.1).

## Results

### Infected ticks were more likely to survive the winter than uninfected ticks

Ticks infected with *B. burgdorferi* had higher overwintering success than uninfected ticks in each of the 3 years (Three-way Loglinear analysis for each year) (Fig. 1). In year 1 (N=36 ticks) the final model demonstrated significant interactions between infection and survival (*X^2^* (1)=4.73, p=0.03) as well as site and survival (*X^2^* (1)=6.1, p=0.01). In year 2 (N=145), the final model demonstrated significant interactions between infection and survival (*X^2^* (1)= 19.6, p<0.001) but not between site and survival (*X^2^* (1)=0.41, p=0.5). In year 3 (N=120) the final model demonstrated a significant interaction between infection and survival (*X^2^* (1)= 5.75. p=0.02), but marginal effects for site and survival (*X^2^* (1)=2.47, p=0.12).

Combining the results from all 3 years, uninfected ticks were more likely to die during the winter than infected ticks (Factorial *X^2^* (1)=24.25, p<0001, N=301 ticks). The overall relative risk of death during overwintering for uninfected ticks (49.25% dead, N=134) vs infected ticks (22.16% dead, N=167) was 1.83 (95% confidence interval 1.4 – 2.4). In other words, uninfected ticks were almost twice as likely to die during the winter as infected ticks.

Tick overwintering mortality varied significantly across years (Year 1: 58.3% (N=21/36, Year 2: 22.1%% (N=32/145), Year 3: 41.7%(N=50/120), post hoc *X^2^* (2)= 21.77, *P*<0.001). The infection rate also varied across years (Year 1: 30.6% (11/36), Year 2: (62.1% (90/145), Year 3: 55.0% (66/120), post hoc *X^2^* (2)=14.8, p=0.0006).

### Temperature and Relative Humidity conditions

The forest sites had higher minimum temperatures than the dune sites (Table S3). Dune grass sites had a greater number of days with minimum temperatures below −5°C, a temperature at which ticks are unlikely to show movement in the field (Clark 1995; Carroll & Kramer 2003). 29.7% days were below −5°C in the dune grass (106 days/357 days over 3 winters) compared with 14.6% of days in the forest sites (52 days/357 days over 3 winters)(*X^2^* (1)=23.7, p<0.0001). Year 1 had the lowest temperatures at the forest and dune grass sites (Table S3), and this year also had the lowest rate of winter survival.

### Implications for tick population growth

Although *B. burgdorferi* does not appear to be vertically transmitted (Rollend et al. 2013), an increase in the survival of infected adult females could lead to an increase in the number of tick larvae in the next generation. Each additional infected female that survives the winter could supply up to 3,000 extra larvae (Mount et al. 1997) to the next generation. After five winters, an area in which 100% of the ticks are infected could have more than 6 times as many ticks as an area with no infected ticks (see Supplementary Information for details).

## Discussion

Overwintering mortality varied across the three years, as has been observed in other field studies (Lindsay et al. 1998). Microclimate had an impact on mortality, but only in the first year in which a higher proportion of ticks survived in the forested site compared to the dune grass site. Other studies have also found that whether ticks have greater overwintering survival in a particular microclimate varies across the years (e.g. Maine (Volk et al. 2022)). The first year was also the coldest year in our study, and it may be that the effect of microclimate is most pronounced during severe winters. Despite this variability in overwintering mortality across years and sites, infected adult female ticks were more likely to survive the winter than uninfected female ticks in each of the three winters at both sites (Fig. 1). Therefore, this effect is robust enough to be detectable across both years and microclimates.

Whether the correlation between infection and winter survival will impact tick population growth depends on whether there is a causal link between infection and overwintering survival. We did not infect our ticks, but collected them from the field. It is possible that the differential survival was caused, not by infection, but by some pre-existing difference(s) between infected and uninfected ticks. For example, it is possible that infected ticks must be in good condition to survive the infection, while in the uninfected ticks, there may be a mix of ticks in good and poor condition. This difference in initial condition between the two groups could explain the difference in host mortality. We argue that this is unlikely. First, *B. burgdorferi* infection produces no measurable damage to ticks and is not pathogenic (Couret et al. 2022). In a related system, *Borrelia afzelii* infection had no effect on the survival of *I. ricinis* nymphs (Hurry et al. 2021). On the contrary, *B. burgdorferi* appears to benefit *I. scapularis* (Couret et al. 2022). Changes in gene expression in infected ticks suggest that infected ticks have an enhanced feeding ability, as well as other advantages (Couret et al. 2022).

Another possibility is that the number of *B. burgdorferi* spirochetes may decrease with time. Therefore, some of our ‘uninfected’ ticks may have been infected ticks that had ‘lost’ their *Borrelia* bacteria due to age. Older ticks are more likely to die, and this would inflate the death rate of uninfected ticks. There is evidence of such a decline in the related tick, *I. ricinis* infected with *Borellia afzelii* (Jacquet et al. 2017). However, even with the decline in bacterial number, there was no change in the detection rate in *I. ricinis* (i.e. qPCR continued to identify *Borrelia* DNA even at the lower bacterial load, Jacquet et al. 2017). In fact, the older the tick, the greater the prevalence of *B. afzelii* in *I. ricinis* (Springer et al. 2022). Perhaps most importantly, the bacterial load of *B. burgdorferi* does not decline in *I. scapularis* nymphs over a 12 month period (see Fig. 2 in Samanta et al. 2022), unlike *I. ricinis* (Jacquet et al. 2017). There may be a difference in the longevity of *Borrelia* in *I. ricinis* compared with its longevity in *I. scapularis*.Finally, *B. burgdorferi* spirochetes have been found to persist in the gut for the entire lifespan of *I. scapularis* (Pal et al. 2021), making it unlikely that they would be ‘lost’ over time. Therefore, there is no evidence that bacterial loss with time would produce a significant false negative rate.

An additional concern is whether it is possible to reliably identify *Borrelia* DNA from dead ticks. We demonstrated that we were able to measure *Borrelia* DNA in ticks that were subjected to daily freeze/thaw for 10 weeks, using a greater temperature fluctuation than they faced in the field (See Supplementary Information). DNA is an extremely stable molecule, with an approximate half-life of 521 years (Allentoft et al. 2012). Moreover, *Borrelia* is found largely in the midgut in the tick (Kurokawa et al. 2020). In humans, the intestinal microbiota of people buried in cold climates from 100 to 3000 years ago can still be identified (Rollo et al. 2007). And although nucleases in the environment do degrade DNA post mortem, low temperatures inactivate these enzymes (Alaeddini et al. 2010) and DNA in dead cells remains intact when frozen (Burkert et al. 2019). For organisms kept at 15°C, it would take 5,000 to 10,000 years for an 800 bp fragment to degrade (Alaeddini et al. 2010). Moreover, the ticks in our study were kept in microcosms, and these tubes isolated them from soil invertebrates and most other organisms, decreasing the risk of degradation. Also, the *Borrelia* DNA was protected by the ticks’ hard exoskeleton during the winter, which was still intact in all of our dead ticks. Perhaps most importantly, we tested the DNA integrity of every sample, and did not find evidence of degradation in the DNA from dead ticks. For these reasons, we do not believe that our results are due to an inability to measure *Borrelia* DNA in dead ticks.

*B. burgdorferi* infection could result in increased winter survival in *I. scapularis* by a number of non-mutually exclusive mechanisms. *B. burgdorferi* infection is known to cause physiological and behavioural changes in *I. scapularis* (Benelli 2020). The evidence suggests that these changes could increase tick condition. For example, *B. burgdorferi* infection enhances feeding in the tick (Courtet et al. 2022), allowing infected ticks to gain more resources during their feeding period. This increase in resources could also result in increased overwintering survival. *I. scapularis* also responds to a *B. burgdorferi* infection with an upregulation of a number of immune genes (Narasimhan et al. 2017; Tang et al. 2021; Hart et al. 2022). In insects, activating an immune response can enhance cold tolerance (Krams et al. 2011). Concomitant with these changes, infection with *B. burgdorferi* induces the expression of a suite of protective genes in *I. scapularis*. For example, *B. burgdorferi* infection results in the increased expression of heat shock proteins (Hart et al. 2022), Glutathione peroxidase and Glutathione-S-transferase (Narasimhan et al. 2017). These type of molecules are known to be cryoprotectant in insects (King & MacRae, 2015; Overgaard & MacMillan, 2017). Therefore, *B. burgdorferi* could enhance winter survival by providing an immune stimulus, leading to the co-activation of a variety of protective mechanisms (e.g. Adamo 2022). We favour this hypothesis, especially as *B. burgdorferi* does not appear to damage ticks (Courtet et al. 2022). In other words, one of the results of the complex interaction between tick and spirochete (Kurokawa et al. 2020) may be hormesis (Berry and Lopez-Martinez 2020), i.e. a mild stressor (i.e. *B. burgdorferi*) leads to improved *I. scapularis* hardiness due to an upregulation of protective mechanisms.

Although *B. burgdorferi* and *I. scapularis* have co-existed for at least 20,000 year (Walter et al. 2017), whether *B. burgdorferi* actively manipulates its host to increase its overwintering survival remains unclear. The ability of an antimicrobial protein to also function as a cryoprotective molecule (Neelakanta et al. 2010) makes the issue difficult to entangle. An additional problem for the manipulation hypothesis, however, is that *B. burgdorferi* in the adult tick has probably reached a dead end in terms of transmission. However, *B. burgdorferi* could benefit by improving host winter survival in larvae or nymphs. Future studies should examine whether infection in these life stages also increases winter survival.

A recent survey of ticks on the south shore of Nova Scotia found that about 4% of ticks were infected with *A. phagocytophilum* (Guillot et al. 2020). Co-infections of *A. phagocytophilum* and *B. burgdorferi* are typically rare, and occur at the rate expected if the pathogens were assorting randomly (Steiner et al. 2008). Therefore, it is unlikely that the enhanced overwintering survival we observed was due to *A. phagocytophilum* infection, not *B. burgdorferi*. Moreover, although our focus was on *B. burgdorferi*, we did run primers for *A. phagocytophilum*, and we did not find it in our samples (see Supplementary Information). Finally, ticks infected with this pathogen could have been in either the *B. burgdorferi* infected or uninfected group. Such a random assortment of *A. phagocytophilum* infected ticks would not produce the large increase in winter survival that we found in ticks infected with *B. burgdorferi*.

Initial tick populations of *I. scapularis* are often *B. burgdorferi* free (Ogden et al. 2010; Ogden et al. 2013). Tick population size needs to reach a certain threshold for *B. burgdorferi* to become endemic (Norman et al. 1999). Our study suggests that infection itself may increase the likelihood of this event by boosting tick population size, especially during winters with low overwintering survival for uninfected ticks.

The potential real world impact of *B. burgdorferi* infection on tick populations urges further study of this phenomenon. If increased overwintering survival of adult females leads to greater population growth, then tick population numbers should grow more rapidly once *B. burgdorferi* infects the population. There is some evidence for this pattern (Elias et al. 2022). But this pattern will only exist where other factors (such as the size of the mouse population, also needed for tick survival) are conducive to a rapid increase in tick numbers (see Ogden et al. 2005; Eisen et al. 2016; Wallace et al. 2019; Ogden et al. 2021; Winter et al. 2021) for a discussion of these factors). Therefore, the true impact of infection on tick numbers will vary from region to region, and, in some cases infection may not provide ticks with an advantage (see discussion in Lemoine et al. 2022).

Regardless of exact nature of the link between *B. burgdorferi* and the increased overwintering survival of infected adult ticks, our results suggest that adult ticks found in the late winter/early spring are more likely to be infected than adult ticks found in the late fall. Unfortunately, we were unable to locate any data to show whether this occurs in areas with endemic *I. scapularis* populations. Whether adult ticks are more likely to be infected with *B. burgdorferi* by mid-spring appears to vary by year (Roome et al. 2018).

Once *B. burgdorferi* enters a population, infection could have a synergistic interaction with other environmental changes (e.g. such as increasing temperatures due to climate change), leading to boosted population numbers in previously marginal areas for *I. scapularis*. In some regions, it may make the difference between the local extinction of a population and its growth.

## Acknowledgements

This research was supported by a Canadian Lyme Foundation Venture Grant (SAA), a Discovery Grant from the Natural Sciences and Engineering Council of Canada (NSERC) (SAA), MITACS Accelerator Award (AEN) and Killam Post Doctoral Fellowship Award (LVF). We thank Dr. L. McMillan and Dr. S. Mackinnon for helpful discussions and thank D. Adamo, A. Barkhouse, L. Mace, and J. Zbarsky for help with DNA extraction and PCR. We thank the Harrison Lewis Discovery Centre for hosting our field study and Thomas Raddall Provincial Park for permission to drag for ticks. We thank the Public Health Agency of Canada (PHAC) for the gift of the OspA for our positive control.

## Disclosure

The authors declare that they have no conflicts of interest to disclose.

## Author Contributions

SAA formulated the hypothesis, assisted with experimental design and data analysis, and led
the writing of the paper. AEN assisted with experimental design and data analysis, wrote a first draft and performed the study. AM validated the qPCR for our study. LVF helped formulate the question, assisted with experimental design and helped write the paper.

## Data Availability

Data are available in the Mendeley repository at: El Nabbout, Amal (2023), “Data 2018-2021 HLCF”, Mendeley Data, V1, doi: 10.17632/tzjdxmyctv.1

## Supplementary Information

### Determination of Tick Infection Status

The procedure used to determine tick infection status was as described in Courtney et al., (2004), and is repeated here. Ticks were identified using standard keys (Keirans and Likwack, 1989). DNA was extracted from *I. scapularis* following the Public Health Agency of Canada (PHAC) protocol, adopted from Courtney et al. (2004). We used a Qiagen QIAamp^©^ DNA Mini Kit with an extra bead tube step (MN Bead Tubes Type D (Macherey-Nagel)) to improve the DNA extraction process. MN Bead Tubes Type D (Macherey-Nagel) were filled with 200 μl of Qiagen ATL buffer. Individual ticks were crushed thoroughly using a Bel-Art^®^ Disposable Pestle. Bead tubes were vortexed for 30 minutes. 20 μl of Qiagen Proteinase K was then pipetted into each bead tube. Bead tubes were incubated at 56°C for 60-90 minutes. 200 μl of Qiagen AL buffer was added to each bead tube and tubes were vortexed for 15 seconds and then incubated at 70°C for 10 minutes. Finally, 200 μl of 100% ethanol was pipetted into each bead tube and then tubes were vortexed for 15 seconds. DNA extraction followed QIAamp^©^ DNA mini kit instructions.

DNA was then screened for evidence of *B. burgdorferi* infection using a multiplex real-time PCR targeting the 23S rRNA of *B. burgdorferi. B. burgdorferi* infection was then confirmed in positive samples using primers targeting the ospA gene. Each sample was tested in triplicates and all plates included: a no template control (5 Microliters of milliQ water), a 25 Cq value positive control, and a 30 Cq value positive control. The thermocycling conditions occurred as follows: activation of enzyme AmpErase at 50°C for 2 minutes, denaturation of AmpErase and activation of AmpliTaq Gold^®^ Polymerase at 95°C for 10 minutes, followed by 40 cycles of amplification at 95°C for 15 seconds and annealing at 60°C for 1 minute.

The integrity of DNA samples was assessed by gel electrophoresis^®^ (Aranda et al., 2012) using a Gel Doc™ XR+ Digital Gel Imaging System^®^ from Bio-Rad. DNA quantity was determined using a Qubit fluorometer.

Although we ran primers for *Anaplasma phagocytophilum*, we found no evidence for it in our ticks, i.e. there was no increase in fluorescence even after 40 cycles.

**Table S1.**
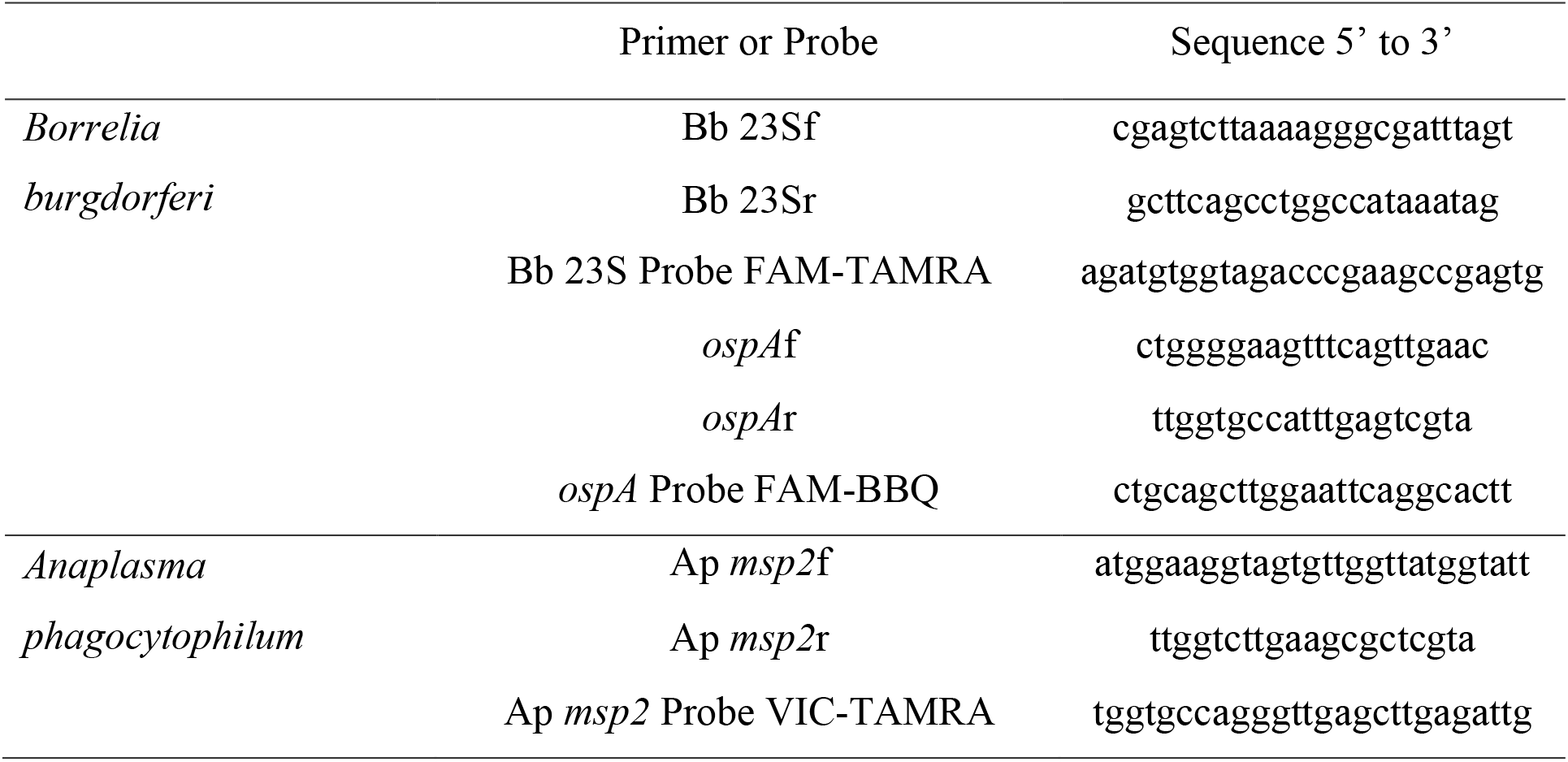
Primer sequences used in Real-time PCR [3].

The efficiency of the Bb23S primer was 92.8%. For the ospA primer, the efficiency was 100.9%.

### Detecting B. burgdorferi in Dead Ticks

Our experimental design depended on our ability to detect *B. burgdorferi* in ticks that may die at various times throughout the field experiment. As a result, we performed experiments to determine whether *B. burgdorferi* would be detectable in dead ticks subjected to freeze/thaw for up to 10 weeks. Ticks were exposed to a fluctuating thermal regime (−5°C to 10°C) in 12-hour increments for 10 weeks This temperature regime exposed these samples to more freeze/thaw events than they would have experienced in the field (Table S3). We focused on the number of freeze/thaw events because DNA in dead cells shows little or no degradation when kept frozen (Burkert et al., 2019), but freeze/thaw cycles accelerate DNA degradation (Rollo et al., 2007). We cut field collected *I. scapularis* ticks in half longitudinally using a sterile razor. The first half was used immediately for DNA extraction. The second half of the tick had a small aliquot of Vaseline added to seal the cut end. This method provided protection from dehydration through the open cut, in the same way that the intact cuticle of a dead tick provides protection from desiccation. These half ticks were stored in a fluctuating thermal regime incubator (−5°C to 10°C) in 12-hour increments for 10 weeks. These temperatures produce more extreme freeze and thaw cycles than was normally observed in the field (Table S3), which assisted us in determining the length of time the DNA is still measurable in *Ixodes scapularis* under Nova Scotian winter conditions. After the 10 week incubation period was completed, we extracted the DNA as previously described and tested for *B. burgdorferi* using qPCR.

Table S2 shows the high concordance (100%) of the results for the two tick halves.

**Table S2.**
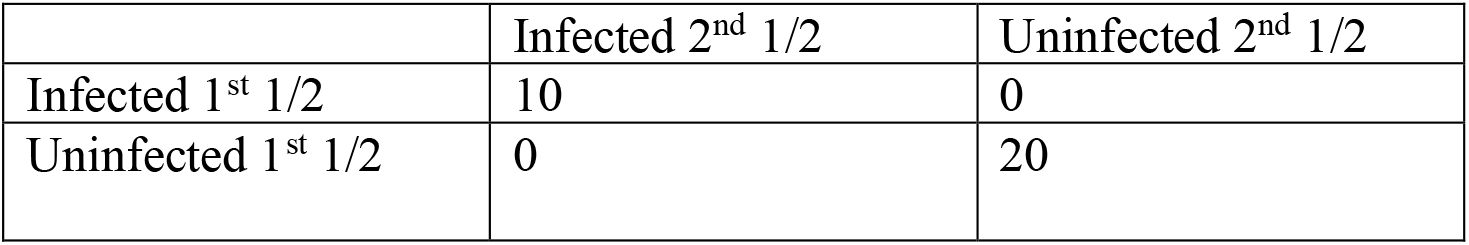
Concordance between the two halves of each tick tested 10 weeks apart.

We also tested the integrity of the DNA from both the lab and field experiments. All DNA was found to be intact, showing distinct bands on the gel (Aranda et al., 2012). These results show that it is possible to reliably determine *B. burgdorferi* infection in the dead ticks we collected.

### Potential Effect of Infection on Tick Population Growth

For our estimate of the potential effect of infection on population growth, we assumed that infection will only change overwintering survival and will not impact other aspects of tick survival (e.g. the likelihood of finding a mammalian host). Using our data, we assumed that infected females have an 80% chance of overwintering survival and uninfected females have a 50% chance of survival. We calculated the relative growth of the two populations, one with 100%*B. burgdorferi* prevalence, another with 0%. We assumed all adult females produced 1,000 larvae and died after laying eggs (Gray and Kahl, 2022). We also assumed that ½ of larvae will be female, and we further assumed that all larvae will survive to overwinter as adult females.

We also assumed that infected adult female ticks pay a 10% fecundity cost, based on data from *Drosophila* (McKean et al. 2008).

---

|  |
|---|
| ◦ |
| ◦ |
| ◦ |
| ◦ |
| ◦ |
| ◦ |
| ◦ |
| ◦ |
| ◦ |

## Notes

### Competing Interest Statement

The authors have declared no competing interest.

### Summary of Updates

Revision highlights important controls.

