## Supplementary material for "Female ticks (*Ixodes scapularis*) infected with *Borrelia burgdorferi* have increased overwintering survival, with implications for tick population growth": Table S3

**Table S3.** Summary of temperature at the forest and dune grass sites over the 2018-2019 (year 1), 2019-2020 (year 2), and 2020-2021 (year 3) winter seasons. Data from year 2 are the average of the two dataloggers at the two different locations at the forest and dune grass sites.

| | Year 1, n=107 days | | Year 2, n=135 days | | | | Year 3, n=115 days | |
|---|---|---|---|---|---|---|---|---|
| | Forest | Dune Grass | Forest 1 | Forest 2 | Dune Grass 1 | Dune Grass 2 | Forest | Dune Grass |
| # of days temperature was <-5°C | 39 | 47 | 3 | 0 | 31 | 18 | 10 | 34 |
| # of days, temperature change >9°C | 19 | 20 | 5 | 4 | 60 | 37 | 9 | 28 |
| Average change in temperature daily, ±STD (°C) | 6.91 ±3.79 | 6.80 2.90 | 4.16 2.35 | 4.05 2.27 | 8.08 3.99 | 7.26 3.50 | 4.58 3.00 | 7.01 3.90 |
| Max daily change in temperature (°C) | 29.4 | 16.1 | 10.8 | 11.6 | 18.85 | 16.3 | 20.85 | 21.15 |
| Min daily change in temperature (°C) | 1.46 | 1.40 | 0.25 | 0.90 | 0.07 | 0.77 | 0.50 | 0.47 |
| Lowest night temperature (°C) | -14.10 | -13.50 | -6.09 | -4.29 | -12.80 | -9.78 | -7.01 | -9.96 |
| Average temperature in the night (°C) | -2.71 | -3.86 | -0.68 | 0.49 | -2.96 | -1.78 | -0.84 | -2.93 |
| Highest daily temperature (°C) | 15.3 | 15.8 | 14.04 | 13.70 | 16.17 | 16.13 | 13.84 | 16.56 |
| Average temperature in the day (°C) | 4.20 | 2.96 | 3.48 | 4.54 | 5.12 | 5.47 | 3.74 | 4.09 |
